# Typing of feces-derived *Candida albicans* strains using a novel seven-locus microsatellite panel reveals associations with yeast phenotype in individuals with inflammatory bowel disease

**DOI:** 10.1101/2024.07.08.601751

**Authors:** Isabelle A.M. van Thiel, Irini A.M. Kreulen, Mèlanie V. Bénard, Marcus C. de Goffau, Bart Theelen, Sigrid E.M. Heinsbroek, Patrycja K. Zylka, Cyriel Y. Ponsioen, Teun Boekhout, Wouter J. de Jonge, Søren Rosendahl, René M. van den Wijngaard, Ferry Hagen

**Author notes:** **Corresponding authors**: Isabelle A.M. van Thiel, PhD. & prof. dr. Ferry Hagen,. Shared second authorship position (Irini A.M. Kreulen, Mèlanie V. Bénard).

## Abstract

Inflammatory diseases of the human gastrointestinal tract are affected by the microbes that reside in the mucosal surfaces along this tract. Patients with inflammatory bowel diseases (IBD) have altered bacterial and fungal intestinal compositions, including higher levels of fecal *Candida* yeasts. Ongoing research indicates that genetic and phenotypic diversity of *Candida albicans* may be linked with disease severity. In this study, we set out to investigate feces-derived *C. albicans* strains from individuals with Crohn’s disease, ulcerative colitis, and healthy volunteers through microsatellite-based genotyping and phenotypic assays. A seven-locus microsatellite panel was applied, of which six loci were newly developed. Combined interpretation of minimum spanning networks and principal component analysis (PCA) led to indications that there is no specific lineage of *C. albicans* that is associated with IBD, but rather that the three study populations do have distinguishable distributions of genotypes. In addition, phenotypic characterization by means of enzyme release assays revealed trends between genotypes and virulence-related enzyme activity. Furthermore, correlations were observed between the clinical inflammation biomarker fecal calprotectin and serum anti- *Saccharomyces cerevisiae* antibodies, as well as between lipase activity of feces-derived *C. albicans* strains and serum 1,3-β-glucan levels. We thus show that microsatellite typing can describe genetic diversity of feces-derived *C. albicans* strains, and that phenotypic diversity of these strains may indeed correlate with fungal genotype or disease. This study opens further possibilities to investigate fecal fungi in relation to severity of inflammation in IBD or in other (intestinal) diseases.

## Introduction

The intestinal fungal community resides in close contact with mucosal surfaces and is thereby able to influence the course and severity of inflammatory bowel diseases (IBD). In the two main subtypes Crohn’s disease (CD)^1^ and ulcerative colitis (UC),^2^ different sections of the gastrointestinal tract can be affected by inflammation. The combined incidence is approximately 0·3% albeit with a Western-predominant pattern,^3^ and symptoms generally include diarrhea, bloody stools, abdominal pain, weight loss, and fatigue.^1, 2^ The pathophysiology of IBD is not fully understood, but several major factors contribute to this disease, including genetics, environmental factors, and the intestinal microbial composition. The latter comprises roughly forty trillion microbes,^4^ of which the vast majority is bacterial, and approximately 0·1 to 1% is estimated to be of fungal origin.^5^ While ample data is available on the role of the bacterial community in IBD, data regarding the gut mycobiome is still limited.

The intestinal fungi composition in IBD patients is generally altered when compared to healthy volunteers, and often characterized by elevated abundances of *Candida* species and lower diversity metrics.^6, 7^ Several murine studies have indicated that colonization of mice with yeasts including *Candida albicans,*^8^ *Debaryomyces hansenii*,^9^ or *Malassezia restricta*^10^ worsen intestinal inflammation. Additionally, genetic mutations in several components of fungal recognition pathways (i.e., *CLEC7A* encoding Dectin-1, *CARD9*) were observed in patients with IBD.^8, 11^ It is hypothesized that this genetic contribution is also reflected by elevated titers of anti-*Saccharomyces cerevisiae* antibodies (ASCAs) in IBD patients, and especially those with CD.^12–15^ These ASCAs recognize yeast mannans and, despite their name, it is assumed that *C. albicans* is the more likely immunological trigger for the presence of ASCAs.^16^ Recent mycobiome studies describe genetic and phenotypic variation of *C. albicans* yeasts that relate to disease severity and mucosal immune responses.^17, 18^ In a study by Li et al., genetic diversity of *C. albicans* strains was determined through whole genome sequencing and subsequently derived single nucleotide polymorphism (SNP) density analyses.^17^ Historically, multiple other methods have been used to map genetic variation, including amplified fragment length polymorphisms fingerprinting (AFLP), random amplified polymorphic DNA (RAPD),^19^ multi-locus sequence typing (MLST) analysis,^20^ and polymorphic microsatellite loci assessment.^21–24^

In the current study, we assessed microsatellite loci to investigate genetic and phenotypic variation of fecal *C. albicans* in the context of CD, UC, and healthy volunteers (HV) as the method provides fast, reliable, and reproducible results.^25^ We developed a novel seven-locus microsatellite panel for *C. albicans* for this purpose. Genetic variation among *C. albicans* strains from one individual was limited, but genetic differences are more pronounced diversity between individuals. A subset of individuals with UC appeared to have a distinct cluster of *C. albicans* strains, even though the study population was very small. In addition, phenotypic evaluations revealed that *C. albicans* lipase activity correlated with detection of 1,3-β-glucans in serum. Taken together, the novel microsatellite panel is useful for assessment of multilocus genotypes and subsequent correlations with phenotypic or clinical parameters in the context of IBD.

## Materials and Methods

### Patients and healthy volunteers

Patients with IBD were recruited at AmsterdamUMC, location University of Amsterdam. For this study, healthy volunteers were defined as “no gastrointestinal abnormalities or disease”. Medication use was allowed in both groups, except for antifungal medications up to three months prior to inclusion. Clinical data was compiled in Castor EDC until analysis of the patient characteristics. Healthy volunteers only provided their sex and year of birth. Medical ethics approval was obtained at AmsterdamUMC, location AMC (number 2017_239). All participants gave written consent for the use of their fecal and blood samples, and patients additionally for the extraction of information from their electronic patient files.

### Sampling and analysis of feces and serum

Patients collected freshly produced fecal samples at home maximally 24 hours before processing. In the meantime, samples were refrigerated. Fecal calprotectin levels were determined on the frozen fecal samples at the routine clinical chemistry laboratory of AmsterdamUMC, location AMC. Blood was drawn at the AmsterdamUMC central phlebotomy laboratory. For determination of ASCAs and 1,3-β-glucans in serum, colorimetric kits were used according to the manufacturer’s protocol, being the Orgentec ASCA IgA/IgG kit (Siemens Healthineers, Den Haag, The Netherlands) and FungiTell diagnostic kit (Nodia, Amstelveen, The Netherlands) respectively. Resulting absorbance of both assays were determined on a Synergy HT plate reader (BioTEK, Beun de Ronde, Abcoude, The Netherlands).

### Culturing and identification of fecal fungi

Small samples of freshly provided stool specimens were diluted using sterile phosphate buffered saline (PBS) and thoroughly vortexed in a 1·5 ml Eppendorf tube. Approximately 100 μl of this suspension was spread onto at least three solid culture media, always including Sabouraud Dextrose Agar with 0·05 g/L chloramphenicol (SAB; Sigma Aldrich, St. Louis, MI, USA), Yeast Peptone Dextrose Agar (YPD; Sigma Aldrich) supplemented with 0·05 g/L chloramphenicol, and malt extract agar (MEA; Oxoid, Basingstoke, United Kingdom) supplemented with 1·68 mg/L penicillin G sodium salt and 1·332 g/L streptomycin sulfate (both Sigma Aldrich). For several samples, Potato Dextrose Agar (PDA), modified Leeming-Notman agar (MLNA) or modified Dixon’s agar (mDA) supplemented with 0·4 g/L cycloheximide and 0·05 g/L chloramphenicol were included as well. Plates were incubated for up to 60 hours at 37°C in an aerobic incubator. When microbial growth was observed, plates were individually sealed and stored at 4°C until further processing.

For identification of yeasts, approximately ten morphologically similar yeast colonies were sub-cultured per individual on glucose yeast peptone agar (GYPA) and subsequently identified in duplicate using MALDI–TOF MS (Bruker Daltronics, Bremen, Germany) using the manufacturer’s protocol for extended direct transfer (eDT). Confirmed yeast strains were routinely frozen in 15% glycerol solutions. Filamentous fungi were omitted from further analysis.

### Yeast DNA extraction

GYPA plates were inoculated with the yeasts of interest and allowed to grow for 16–48 hours. Biomass was collected using a sterile culture loop, and samples were frozen until extraction of DNA, which was performed using the Wizard Genomic DNA Purification kit (Promega, Leiden, The Netherlands). In brief, yeast cells were mechanically disrupted using glass beads in Nucleic Lysis Solution. After a 30–minute incubation period at 65°C, suspensions were subjected to RNase A treatment (Promega) for 15 minutes at 37°C. Protein Precipitation Solution (Promega) was added to remove proteins from the solutions, and DNA was lastly precipitated in isopropyl alcohol. After washing of the pellet in 70% ethanol, DNA was dissolved in DNA Rehydration Buffer (Promega). Presence and integrity of DNA were confirmed using 1·5% agarose Tris-Acetate-EDTA (TAE) gel electrophoresis.

### Bioinformatic analysis of STR lengths

The microsatellite dataset was converted into a *Genind object* and analyzed using the packages adegenet^26^ v2.1.7 and poppr^27^ v.2.9.3 in R v.4.2.1. A matrix based on Bruvo’s genetic distances,^28^ was constructed and used to generate a minimum spanning tree based using the function *plot_poppr_msn* in poppr. A discriminant analysis of principal components was run using the adegenet package 2.1.7 (updated 2022) in R. DAPC analyses were conducted with *de novo* grouping without considering the three patients groups. We used the *find.clusters()* function to determine the number of groups (K) de novo. The optimal number of PCs to use in the DAPC was determined using the optim.a.score() and xvalDapc() commands and 1000 replicates.

### Microsatellite typing

Short tandem repeat (STR) lengths were determined for seven loci of cultured obtained *C. albicans* yeasts. One of these loci, CAI, was previously described.^29, 30^ Primers for the remaining six loci were designed using Tandem Repeat Finder^31^ based on the genome of *C. albicans* SC5314 (=CBS 8758). After selecting suitable repeats, primers on flanking regions were designed using Primer3 v0.4.0 (Table 1).^32^ Forward primers were labeled with 6-carboxyfluorescein on the 5’-terminus. A BioTaq polymerase kit was used for all PCR reactions, which consisted of 1× NH_4_ reaction buffer, 1·5 mM MgCl_2_, 0·04 μM dNTPs, 0·5 U BioTaq (all Bioline, Meridian Bioscience, Cincinnati, OH, USA) and 1 pmol of each primer (Integrated DNA Technologies (IDT), San Diego, CA, USA). Amplification was performed according to the following program: 5 minutes at 94°C, followed by 35 cycles of 30 seconds at 94°C, 30 seconds at 60°C, and 60 seconds at 72°C. Final elongations were performed at 72°C for 5 minutes. The resulting amplicons were purified using Sephadex G–50 and diluted 200–fold in water. Hereof, 1 μl was mixed with 8·9 µl water and 0·1 μl Orange 600 DNA size standard (Nimagen, Nijmegen, The Netherlands) followed by incubation at 100°C for 1 minute.

**Table 1.**
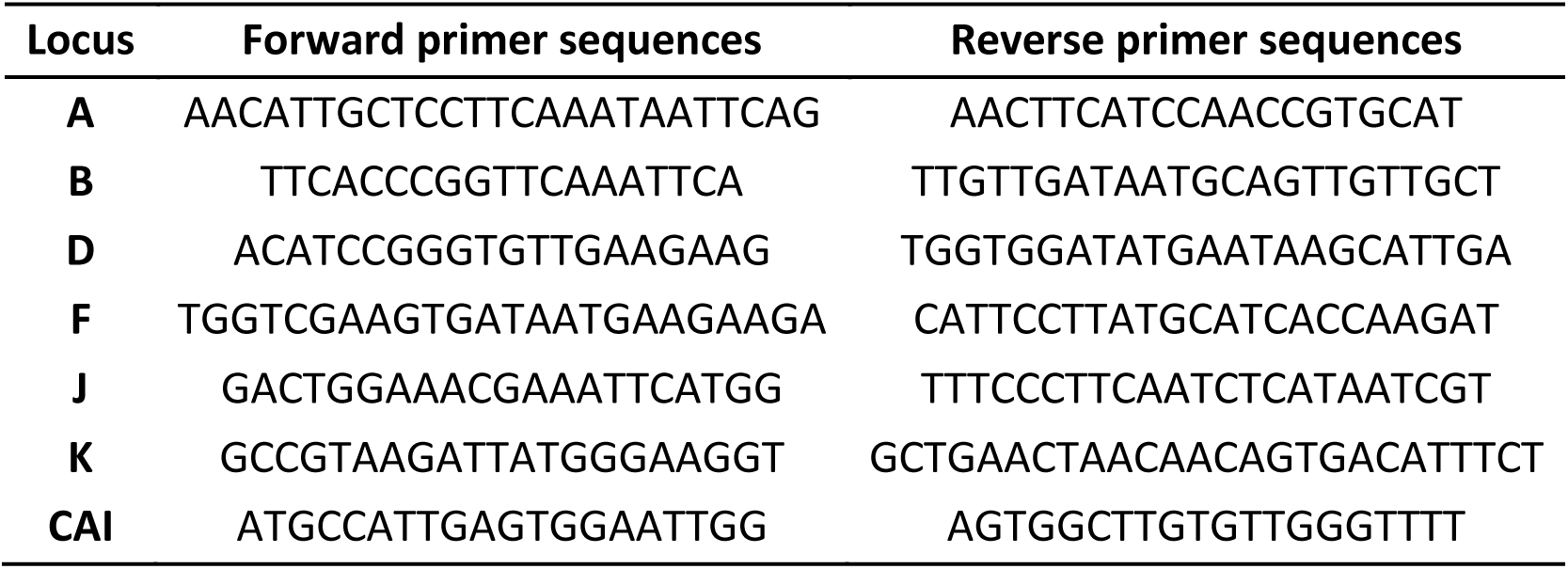
Primer sequences microsatellite loci. Forward primer sequences are labeled with 6–carboxyfluorescein on the 5’-terminus.

Fragment lengths were determined by capillary electrophoresis on an ABI3730xL Genetic Analyzer platform (Applied Biosystems, Palo Alto, CA, USA). BioNumerics v7.6 (Applied Maths, Sint-Martens-Latem, Belgium) was used for peak processing and generation of minimum spanning trees. Inability to read short tandem repeat (STR) lengths for more than two loci led to exclusion of the sample from the analysis.

### Determination of enzymatic activity

Virulence-related enzyme activity of *C. albicans* strains was determined using multiple solid substrate media as previously described.^24^ Overnight *C. albicans* cultures (30°C, 200 rpm) in Yeast Extract Peptone Dextrose Broth (YPD; 1% Yeast Extract, 2% Peptone, 2% Dextrose) were used. Cultures were washed twice in Dulbecco’s phosphate buffered saline (dPBS) before counting using a Nexcelom Cellometer K2 cell counter (Nexcelom Bioscience, Manchester, United Kingdom) and adjusting the inoculum to 1×10^8^ cells/ml in dPBS. A drop of 3 μl of this suspension was spotted in triplicate onto one of the specified culture media. For lipase activity, tributyrin agar was used.^33^ Egg yolk-containing Sabouraud Dextrose agar was used to determine phospholipase activity.^34^ Tween-80 containing agar served as medium for release of esterase.^35^ For proteinase activity, bovine serum albumin (BSA) was added to the agar medium.^36^ Plates were placed at 37°C for varying incubation times: phospholipase activity was determined after 3 days, proteinase and esterase after 5 days, and lipase activity after 6 days of incubation. Activity of the released enzymes was calculated as 1 – (diameter colony/diameter precipitation zone). Experiments were performed in three independent replicates and the *C. albicans* strain SC5314 served as reference.

### Statistical analysis

Baseline characteristics of included individuals were summarized in R v4.1.1 using package tableone v0.13.0. Characteristics were, in line with STROBE (Strengthening the Reporting of Observational Studies in Epidemiology) guidelines for cross-sectional studies, not tested for significance.^37^ *In vitro* data regarding *C. albicans* enzyme release is displayed as mean and standard deviation. Statistical differences between groups were tested using Mann-Whitney U test or Kruskal–Wallis test and Dunn’s multiple comparisons test. Differences in enzyme activity *versus* the activity of the reference strain were determined using a one-sample t–test. Correlations were based on Spearman r. *p* values below 0·05 were considered statistically significant. Visualization of PCA plots and analysis of *in vitro* data were performed in GraphPad Prism v9.5.1.

## Results

### Culturing of fecal samples predominantly results in *Candida* spp

A total of 46 patients (*n*=33 CD*, n*=12 UC, *n*=1 IBD-unclassified) were recruited at the AmsterdamUMC and 17 healthy volunteers (HV) were enrolled (Table 2, Supplementary Table 1) between January 2018 and July 2021. The majority of IBD patients (65·2%) was in clinical remission at the time of inclusion following physician assessment of the electronic patient files. Fecal cultures resulted in growth of any yeast in 72·6% of the cases (CD 36·4%, UC 75·0%, HV 70·6%; Table 3). Among these, species of the *Candida* genus were most frequently observed throughout all groups (CD 66·7%, UC 58·3%, HV 58·8%), and the number of *C. albicans* positive cultures was higher for UC patients compared to CD and healthy volunteers (CD 42·2%, UC 75·0%, HV 53·0%).

**Table 2.** Patient characteristics. All data are expressed as mean (standard deviation) or number of observations (%). BMI, body mass index; EPF, electronic patient file. *Biologicals include anti-TNF, interleukin inhibitors, vedolizumab, and JAK inhibitors. Clinical remission based on electronic patient file is physician guided. Number of observations: fecal calprotectin, *n*=60; Disease duration, *n*=44; BMI, *n*=34; HBI, *n*=18; SCCAI *n*=8. For Montreal disease severity classification, see Supplementary Table 1.

| *n* |  | CD<br>33 | UC<br>12 | HV<br>17 |
| --- | --- | --- | --- | --- |
| Female (%) |  | 21 (63.6) | 5 (41.7) | 13 (76.5) |
| Fecal calprotectin ($\mu\text{g/g}$ feces) | | 281.03 (525.14) | 210.83 (575.09) | 23.00 (23.53) |
| BMI ($\text{kg/m}^2$) | | 25.19 (4.48) | 25.41 (2.82) | |
| Duration of disease (years) |  | 14.90 (9.17) | 16.25 (10.66) |  |
| Medication usa (%) | No IBD medication | 1 (3.0) | 0 (0.0) |  |
|  | Aminosalicylates | 3 (9.1) | 7 (58.3) |  |
|  | Thiopurines | 12 (36.4) | 7 (58.3) |  |
|  | Steroids | 4 (12.1) | 0 (0.0) |  |
|  | Biologicals or JAK inhibitors | 18 (54.5) | 6 (50.0) |  |
|  | Other IBD medications | 3 (9.1) | 2 (16.7) |  |
|  | Proton pump inhibitors | 5 (15.2) | 4 (33.3) |  |
| Intestinal resection (%) |  | 1 (3.0) | 0 (0.0) |  |
| Clinical remission based on EPF (%) | Yes | 19 (57.6) | 11 (91.7) |  |
|  | No | 10 (30.3) | 1 (8.3) |  |
|  | Unclear | 3 (9.1) | 0 (0.0) |  |
| Harvey-Bradshaw Index |  | 2.67 (3.38) |  |  |
| Simple Clinical Colitis Activity Index |  |  | 1.50 (1.60) |  |

**Table 3.**
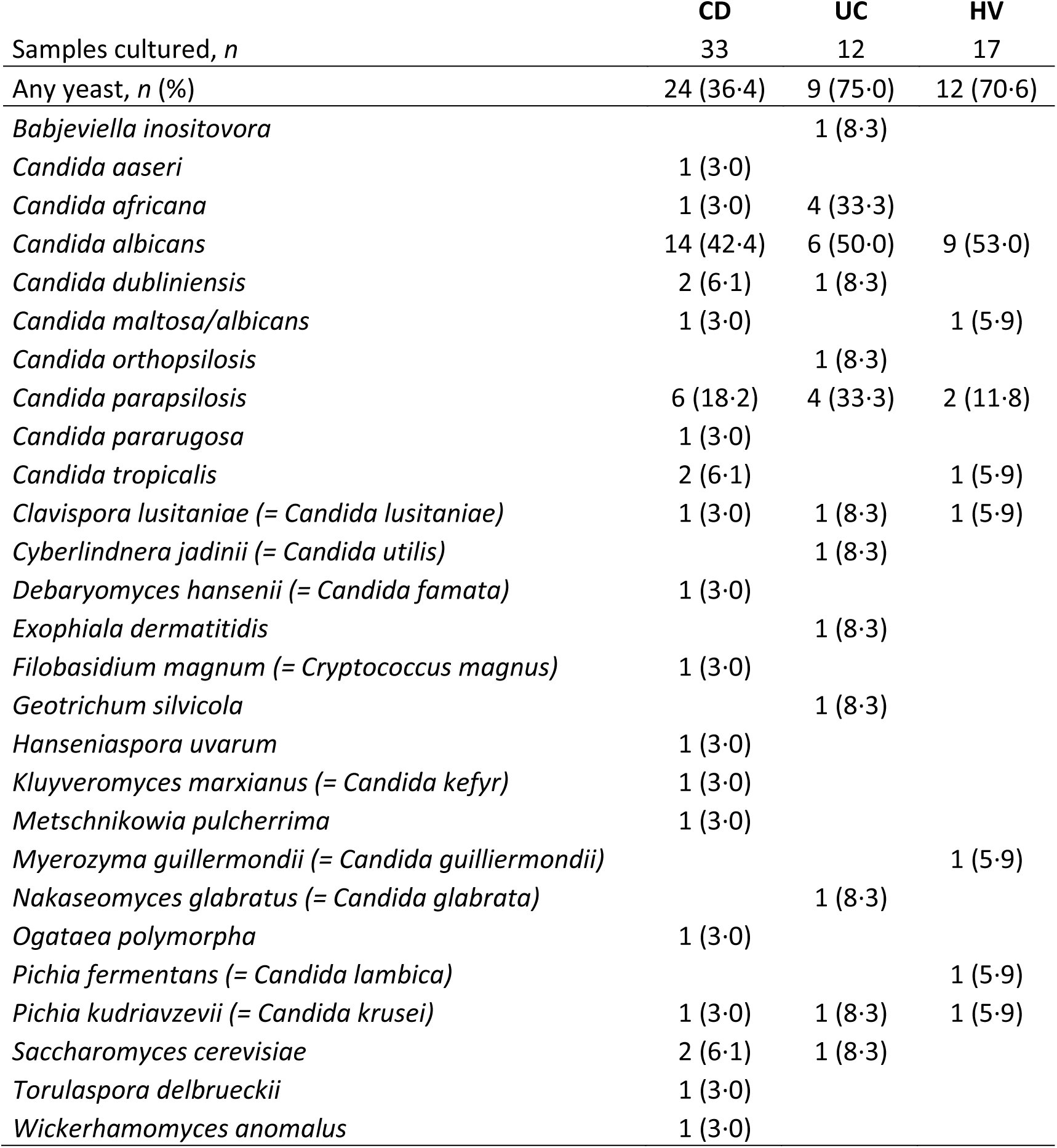
Cultured fungal species from fecal samples. Table indicates number of samples positive for each fungal species detected and percentage of total cultures in brackets. CD, Crohn’s disease; UC, ulcerative colitis; HV, healthy volunteer.

### Genetic evaluation of *C. albicans* strains shows inter- and intra-individual variation

Given earlier reports on genetic variation of *C. albicans* yeasts in patients with IBD and healthy volunteers,^17, 21–24, 38^ we performed genetic analysis of the obtained *C. albicans* strains. To this end, a novel panel of seven microsatellite loci was developed. For six loci, primers were newly developed and not in particular relation to any gene, and the seventh marker was the previously described locus CAI.^29, 30^ A minimum spanning network (MSN) was generated based on the resulting STR analyses of 257 strains derived from 29 individuals (Figure 1A). In total, 77 multilocus genotypes of *C. albicans* were observed. Genetic variation mainly occurred between individuals rather than within individuals (Supplementary Figure 1A-B). Yet, visually striking, no UC strains are found in the lower right branch of the MSN Figure 1A. In addition, it appears that patients with UC relatively have fewer genotypes per individual (Figure 1B), albeit not statistically significant in the current cohort. No statistical conclusions can thus be drawn with respect to the association between *C. albicans* genotype and disease using the microsatellite-based MSN analyses.

**Figure 1.**
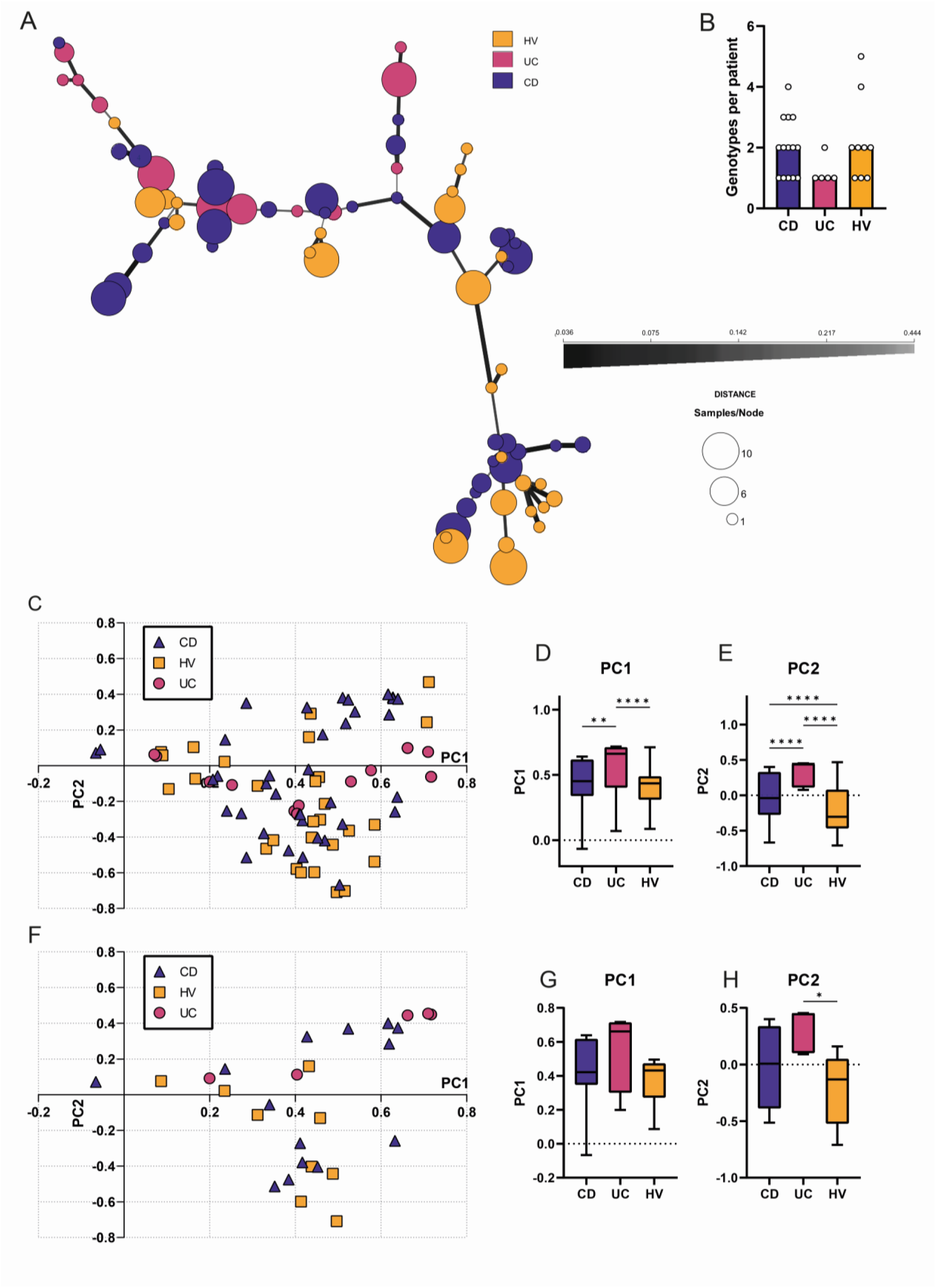
Analysis of genetic diversity of feces-derived *Candida albicans* strains through minimum spanning networks and principle component analysis. N=29 individuals (*n*=14 CD, *n*=5 UC, *n*=9 HV) were included in this analysis, resulting in *n*=257 *Candida albicans* strains. MST was determined based on seven microsatellite loci and is not clone corrected. Thickness and style of the line indicates similarity between different clusters based on Bruvo’s distance. B) Number of genotypes observed per patient. **C**) PCA analysis of all included *Candida albicans* strains mainly shows separation of diseases along PC2 rather than PC1. D-E) Quantification of principle component scores along PC1 and PC2. Ulcerative colitis (UC) has a distinct position along PC1 and PC2 compared with Crohn’s disease (CD) and healthy volunteers (HV).) PCA based on corrected data in which the ‘median’ clone per patient is shown. Kruskal-Wallis test, * p<0·05, ** p<0·01, **** p<0·001.

### Principal component analysis of microsatellite data reveals separation of UC-derived *C. albicans* strains

A principal component analysis (PCA) was additionally performed to statistically differentiate the three study groups from one another. In order to perform this PCA, the STR length of each of the alleles of the seven loci was binarized. For each of the 14 measurements (7 loci × 2 alleles), a sample can thus only be positive for one length option (Supplementary File 1). This analysis excludes any assumptions on genetic relatedness between the different length options. Based on this analysis, a statistically significant separate cluster of UC-derived *C. albicans* strains can be observed, both along PC1 and PC2 (Figure 1C-E), and all study populations have a significantly different distinct distribution along PC2 (Figure 1E). However, as the inclusion of multiple strains per individual may skew statistical analysis, the same analysis was repeated but with only one strain per individual, in which the most abundant strain per individual was selected as representative, or in case of a tie, the median coordinates of all strains of an individual was used instead (Figure 1F-H). Despite the loss of power, the distinct position of *C. albicans* strains from patients with UC on PC2 remained significant in comparison to healthy volunteers (Figure 1G-H). In contrast, discriminant analyses of principle components (DAPC) analysis is specifically able to determine relations between (possible) different clusters by optimizing variance between groups while minimizing within group variance.^39^ The clone-corrected data was subjected to DAPC analysis, which resulted in four clusters (Supplementary Figure 2A). The three disease groups were scattered across the four clusters, although UC-derived strains were not observed in clusters 1 and 3 (Supplementary Figure 2B). Thus, it appears that *C. albicans* strains from UC patients have a distinct genotype compared to healthy volunteers and patients with CD, represented by scoring higher on average on PC2.

### Clinical parameters and enzyme activity of feces-derived *C. albicans* strains correlate

To investigate relations between genotype and phenotype of the feces-derived *C. albicans* strains, we proceeded with phenotypic investigation as described by means of enzyme release for a single strain per individual.^24^ Activities for four enzyme groups were determined, being proteinases, phospholipases, lipases, and esterases (Figures 2A-E, Supplementary Figure 2). The enzyme activity levels of some individual strains deviated significantly from the reference strain SC5314 (Supplementary Figure 2A-D), but when all strains are compiled into their respective study populations, no statistically significant differences were established (Figure 2A-E). Nevertheless, proteinase activity was slightly higher in UC-derived strains (Figure 2A), and esterase activity is potentially slightly elevated in CD-derived *C. albicans* strains (Figure 2D). We additionally examined clinical markers in serum samples of all included individuals (Supplementary Figure 3). Fecal calprotectin, a commonly used clinical marker for intestinal inflammation, was only elevated in the CD group compared with healthy volunteers (Figure 2F). ASCA titers (IgA, IgG, and total) were also significantly altered between the total groups of CD and HV, although this statistically significant difference disappeared when the analysis was only focused on individuals of whom fecal culture was positive for *C. albicans* due to the reduced group sizes (Figures 2F-J). In general, only marginal differences were observed in regards to biomarkers and the three study groups.

**Figure 2.**
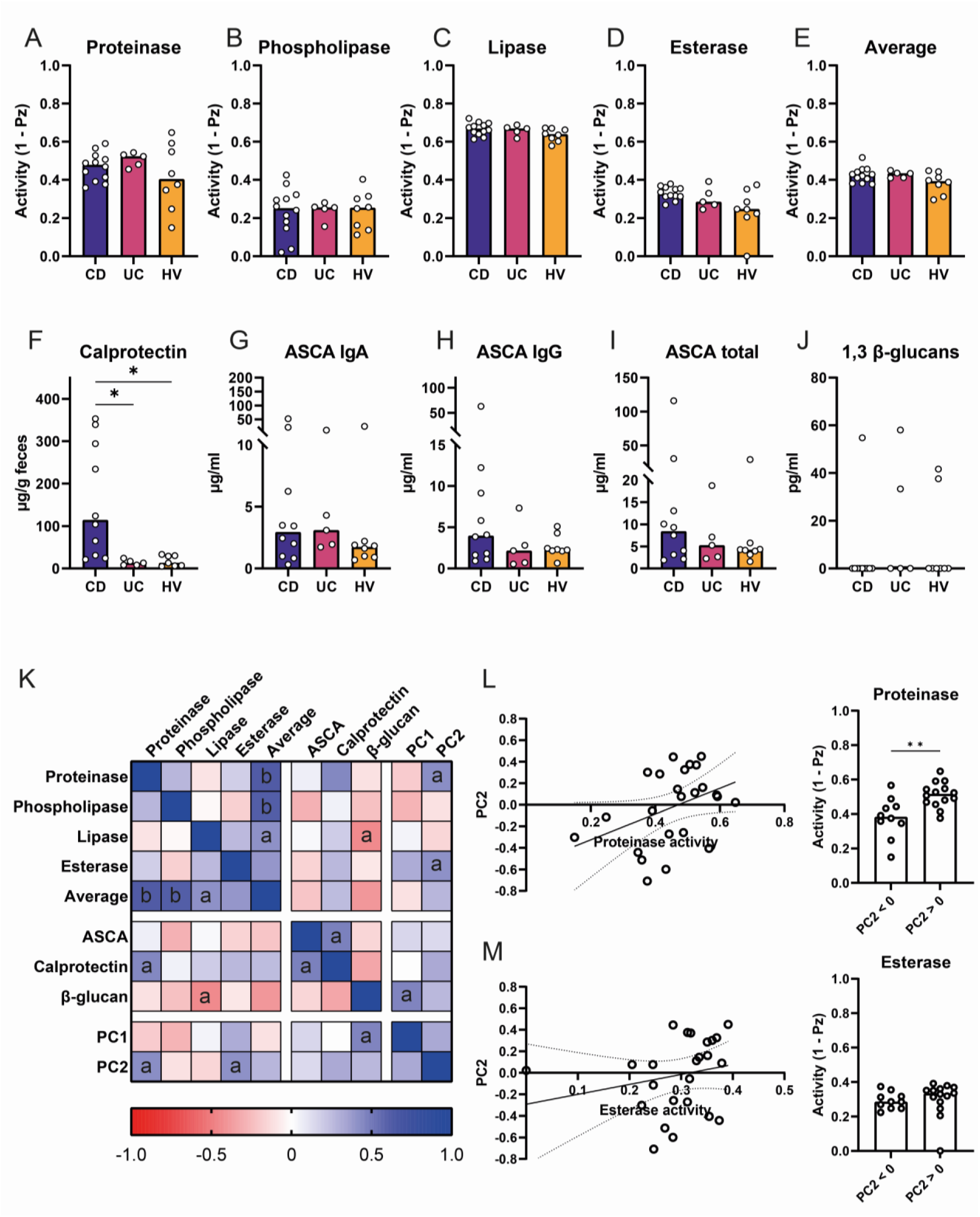
Enzymatic activity of feces-derived *Candida albicans* strains reveals no differentially active enzymes. A-E) Enzyme activity was based in triplicate for one strain per individual. Each data point represents mean activity of a strain as determined in triplicate. A) Proteinase activity. B) Phospholipase activity. C) Lipase activity. D) Esterase activity. E) Average activity across all enzymes. Horizontal dashed line indicates activity of reference strain SC5314. F-J) Clinical fecal and serum titers. F) Fecal calprotectin. G) Anti-*Saccharomyces cerevisiae* antibody (ASCA) titers, IgA immunoglobulins. H) ASCA titers, IgG immunoglobulins. I) Total ASCA titers. J) Serum 1,3-β-glucan levels. K) Correlation plot of all determined enzyme activities and clinical parameters based on Spearman r. Letters and symbols denote significance levels. Spearman r, ^a^ p <0·05, ^b^ p<0·005. L) Correlation between PC2 and proteinase activity. Spearman r=0·417, *p*=0·043, *n*=24. M) Correlation between esterase activity and PC2. Spearman r=0·408, *p*=0·048, *n*=24. Data of all included patients, except for enzyme activity (*n*=8 HV, *n*=12 Crohn’s disease, n=4 Ulcerative colitis). Kruskal-Wallis or Mann-Whitney U test, * p<0·05, ** p<0·01

As previous research has shown that phenotypic characteristics of *C. albicans* strains may correlate with disease severity, Spearman correlation coefficients were determined to investigate relations between each of the acquired parameters (Figure 2K). Of interest is that significant correlations were observed between fecal calprotectin levels and (total) ASCA levels (r=0·464, *p*=0.04, *n*=20), lipase activity of feces-derived *C. albicans* strains and serum 1,3-β-glucan levels (r=-0·448, *p*=0.036, *n*=21). Lastly, significant trends were observed between fecal calprotectin and proteinase (r=0·441, *p*=0.046, *n*=21). Thus, although the enzymatic activities are not significantly higher or lower in any of the three study groups, several correlations exist between clinical parameters and enzymatic activities.

### C. albicans enzymatic activity correlates with PCA-based genetic separation

When linking phenotypic characteristics with PCA coordinate localization various other relevant differences emerge. A strong correlation was observed between PC2 and proteinase activity (Figure 2K-L; r=0·417, *p*=0·043, *n*=24) and esterase activity (Figure 2M; r=0·408, *p*=0·048, *n*=24) highlighting that the lack of significance in regard to UC and the above disease parameters is likely a power issue. For the DAPC analysis, no clear trends were found between genetic cluster and enzyme activity or clinical parameters, with the exception of a possible link between DAPC cluster and ASCA IgA titers (Supplementary Figure 2C-L). In conclusion, genetic and phenotypic data modalities of feces-derived *C. albicans* strains were shown to be linked through PCA-analysis.

## Discussion

Genetic and functional differences among the intestinal fungal populations may contribute to intestinal inflammation. Earlier research already described a potential role for *C. albicans* in the inflammatory bowel diseases UC and CD, ^8, 11, 14, 15^ and the most recent studies indicate that differences at the strain level were related to disease severity and clinical parameters like serum ASCA titers.^17, 18^ In the current study, we focused on feces-derived *C. albicans* strains from patients with either UC or CD, and included a cohort of control individuals. Using a newly developed microsatellite panel of seven loci, we investigated the genetic diversity of *C. albicans* strains within and between individuals using minimum spanning network and PCA analyses. PCA analyses revealed that the strains derived from UC patients tend to be more alike. Attempting to correlate genetic and functional diversity, one strain per individual was subjected to phenotypic examination by means of enzymatic activity determination. Correlations between enzymatic activity and clinical markers were observed, as well as associations between genetic distribution by PCA and enzymatic markers. Taken together, we show that our novel microsatellite panel is able to discriminate feces-derived *C. albicans* strains and that functional capacities may contribute to intestinal disease.

The intestinal tract harbors many different fungal species, but which ones of these are viable in the challenging and competitive intestinal environment is not fully elucidated yet. Culturing feces-derived fungi remains challenging due to the wide variety in nutritional and atmospheric requirements. In our current study, we cultured fecal samples of 62 individuals under aerobic conditions on various culture media, resulting in 27 different yeast species. Since *C. albicans* was the most frequently observed yeast in the fecal cultures, we proceeded our investigations with this species. Earlier reports of *C. albicans* in IBD mainly reported high abundances or overgrowth of this yeast in fecal samples of patients at the species level, and similar observations have been made through ITS1-based metabarcoding on fecal or mucosal samples. Because recent studies indicated that genetic and functional sub-strain differences could contribute to inflammation, we have focused solely on qualitative, culture-based methods. Thus, whether the patients included in this study have increased abundances of *C. albicans* was not determined. Given the large number of strains, we were able to perform genetic analysis on the obtained specimens. Microsatellite analysis provides a relatively rapid and reproducible manner of assessing genetic variation.^25^ As such, we were able to map genetic diversity between and among individuals, either with an inflammatory bowel disease or for healthy subjects. Earlier, similar analyses were performed on feces-derived *C. albicans* strains using amplified fragment length polymorphism fingerprinting (AFLP), random amplified polymorphic DNA (RAPD) analysis, and whole-genome sequencing.^17, 19, 20, 24, 40^ Across all studies, a similar pattern arises which describes mainly inter-individual genetic variation. Therefore, the microsatellite panel developed for this study is a useful tool for genotyping of feces-derived *C. albicans* strains.

Genetic and phenotypic diversity of UC-derived fecal *C. albicans* was previously also described by Li et al.^17^ Genetic diversity was assessed through two methods, being whole genome sequencing and subsequent SNP density analyses. No immediate correlation between disease status and genotype was observed, although a separation of patient-derived strains was visible based on how aggressively the *C. albicans* strains behaved towards murine bone-marrow derived macrophages. The damage-inducing potential of *C. albicans* was also translated to an *in vivo* experimental setting, where the high-damage strain would induce more severe inflammation.^17^ In our current study, genetic differences were observed between UC and CD or HV derived *C*. *albicans* strains, which has not yet been described before. However, based on our results, we cannot yet conclude whether the obtained strains would also be able to differentially influence disease in the human (inflammatory) setting. The *in vivo* situation of IBD is highly complex and influenced by multiple factors, including the (interplay between) bacterial and fungal microbial community, as well as genetic variations within the host that may influence immunological responses.^41–43^ Thus, although we observed genetic separation of UC-derived *C. albicans* strains, further studies should investigate whether a causal relation exists between this and disease (severity).

In conclusion, we here present a novel seven-locus microsatellite typing panel which can describe inter- and intra-individual genetic variation among feces-derived *C. albicans* strains. PCA-based analysis, and visually the analysis of minimum spanning networks, show separation and clustering of ulcerative colitis-derived fungal strains. Phenotypic analysis through assessment of enzyme release shows correlations with genotype and clinical markers, although not significantly with the disease subtype. Together, this research opens the possibility to further study yeasts in (inflammatory) bowel diseases through microsatellite locus assessment.

## Author contributions

Conceptualization, S.E.M.H., T.B., R.M.v.d.W., and F.H.; Methodology, I.A.M.v.T, M.C.d.G., B.T., S.E.M.H., T.B., S.R., R.M.v.d.W., and F.H.; Formal analysis, I.A.M.v.T., I.A.M.K., M.C.d.G., and S.R.; Investigation, I.A.M.v.T., I.A.M.K., M.V.B., and P.K.Z.; Resources, M.V.B. and C.Y.P.; Data curation, I.A.M.v.T., B.T., and F.H.; Writing original draft, I.A.M.v.T.; Writing – Review & Editing, all authors; Visualization, I.A.M.v.T., I.A.M.K., M.C.d.G, P.K.Z., and S.R.; Supervision, T.B., R.M.v.d.W., and F.H.; Project administration, I.A.M.v.T., T.B., and F.H.; Funding acquisition, S.E.M.H, W.J.d.J., T.B., R.M.v.d.W., and F.H.

## Data availability

Medical data will not be made available due to privacy regulations. Data generated in this study is available in the Supplementary Materials upon publication of this manuscript.

