## Supplemental Table and Figures for "Typing of feces-derived *Candida albicans* strains using a novel seven-locus microsatellite panel reveals associations with yeast phenotype in individuals with inflammatory bowel disease"

### Supplementary table and figures

**Supplementary Table 1 | Extended baseline characteristics, Montreal classification.** Classifications Age, Location, and Behavior are described for Crohn’s disease (CD) only, and Extensity and Severity are for description of ulcerative colitis (UC). GI, gastrointestinal. Number of observations: Age, behavior, *n*=33; Location, *n*=46; Severity, *n*=12; Extensity, *n*=11.

|  |  | **CD** | **UC** |
| --- | --- | --- | --- |
| *n* |  | 33 | 12 |
| **Age at diagnosis (%)** | A1: <17 years | 3 (9·1) |  |
|  | A2: 17-40 years | 23 (69·7) |  |
|  | A3: > 40 years | 7 (21·2) |  |
| **Location (%)** | L1: terminal ileum | 9 (27·3) |  |
|  | L2: colon | 13 (39·4) |  |
|  | L3: ileocolon | 11 (33·3) |  |
|  | L4: upper GI disease | 2 (6·1) |  |
| **Behavior (%)** | B1: non-stricturing, non-penetrating | 25 (75·8) |  |
|  | B2: stricturing | 4 (12·1) |  |
|  | B3: penetrating | 4 (12·1) |  |
|  | P: perianal disease | 5 (15·2) |  |
| **Extent (%)** | E1: proctitis |  |  |
|  | E2: left-sided UC |  | 6 (50·0) |
|  | E3: extensive UC |  | 5 (41·7) |
| **Severity (%)** | S0: remission |  | 11 (91·7) |
|  | S1: mild symptoms |  |  |
|  | S2: moderate symptoms |  |  |
|  | S3: severe symptoms |  | 1 (8·3) |

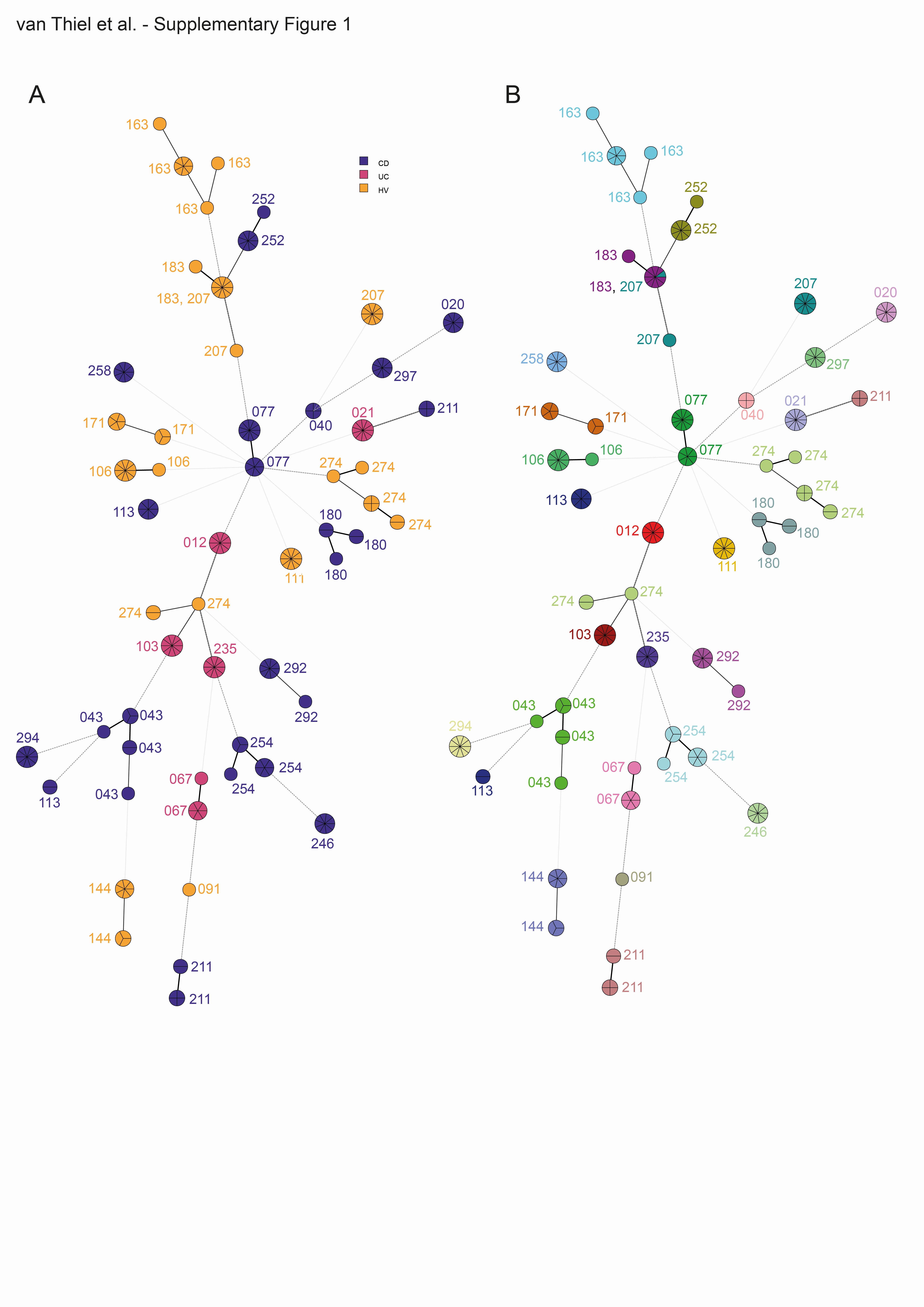
**Supplementary Figure 1 | Minimum spanning trees of all included *C. albicans* strains.** A) Nodes of the minimum spanning tree colored by diagnosis. B) Minimum spanning tree in which the nodes are colored by patient ID. Length and width of lines correspond with the degree of relatedness between nodes.

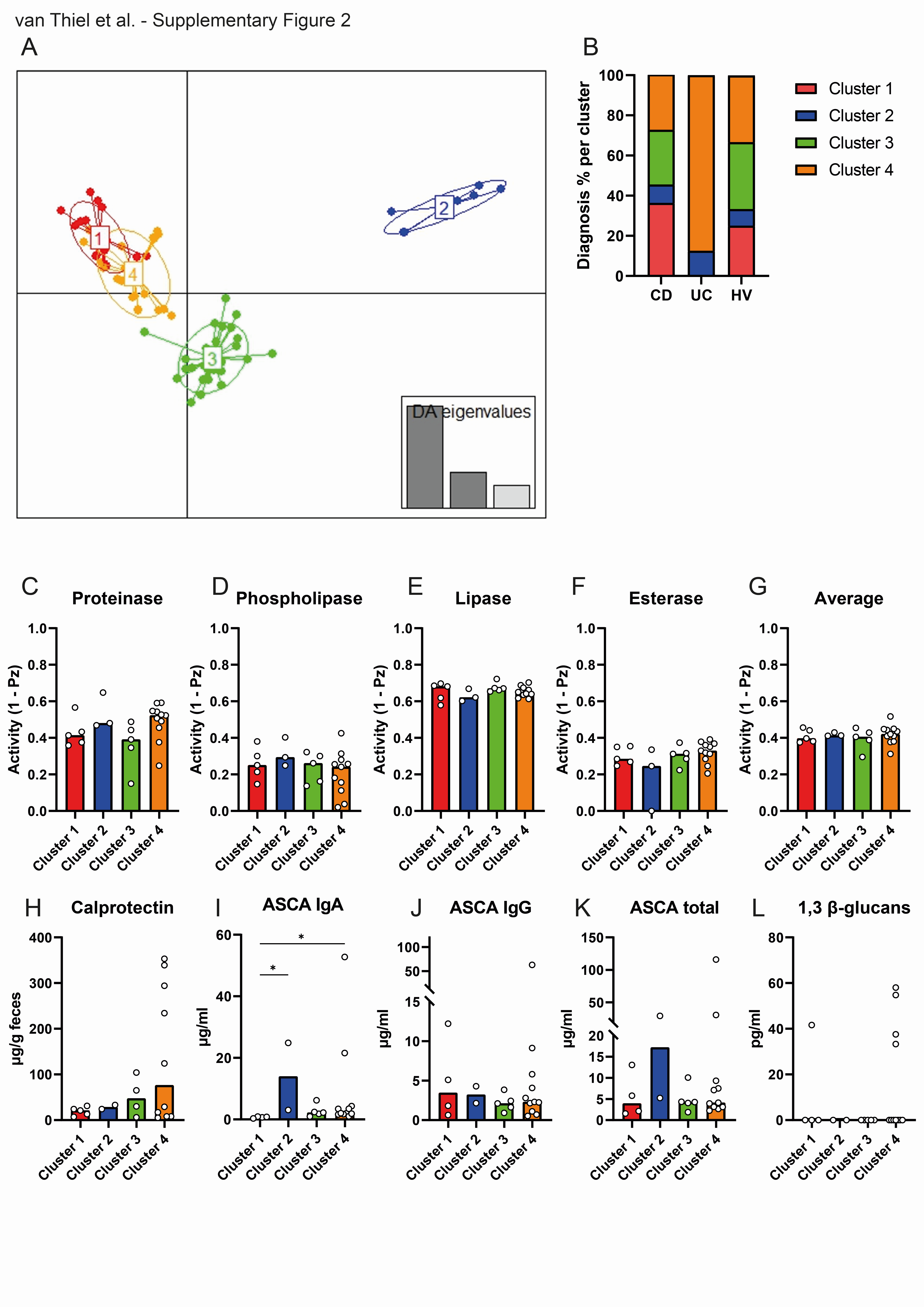

**Supplementary Figure 2 | Extended *Candida albicans* population analyses based on microsatellite typing on seven loci reveals clustering between disease phenotypes.** A) DAPC analysis of clone-corrected *Candida albicans* strains results in four clusters. B) Assignment of strains across clusters based on the disease diagnosis of the individual. C-L) Phenotypic and clinical data as presented in Figure 3, displayed per assigned cluster. Horizontal dashed line indicated activity of reference strain SC5314. Kruskal-Wallis test, * p<0.05.

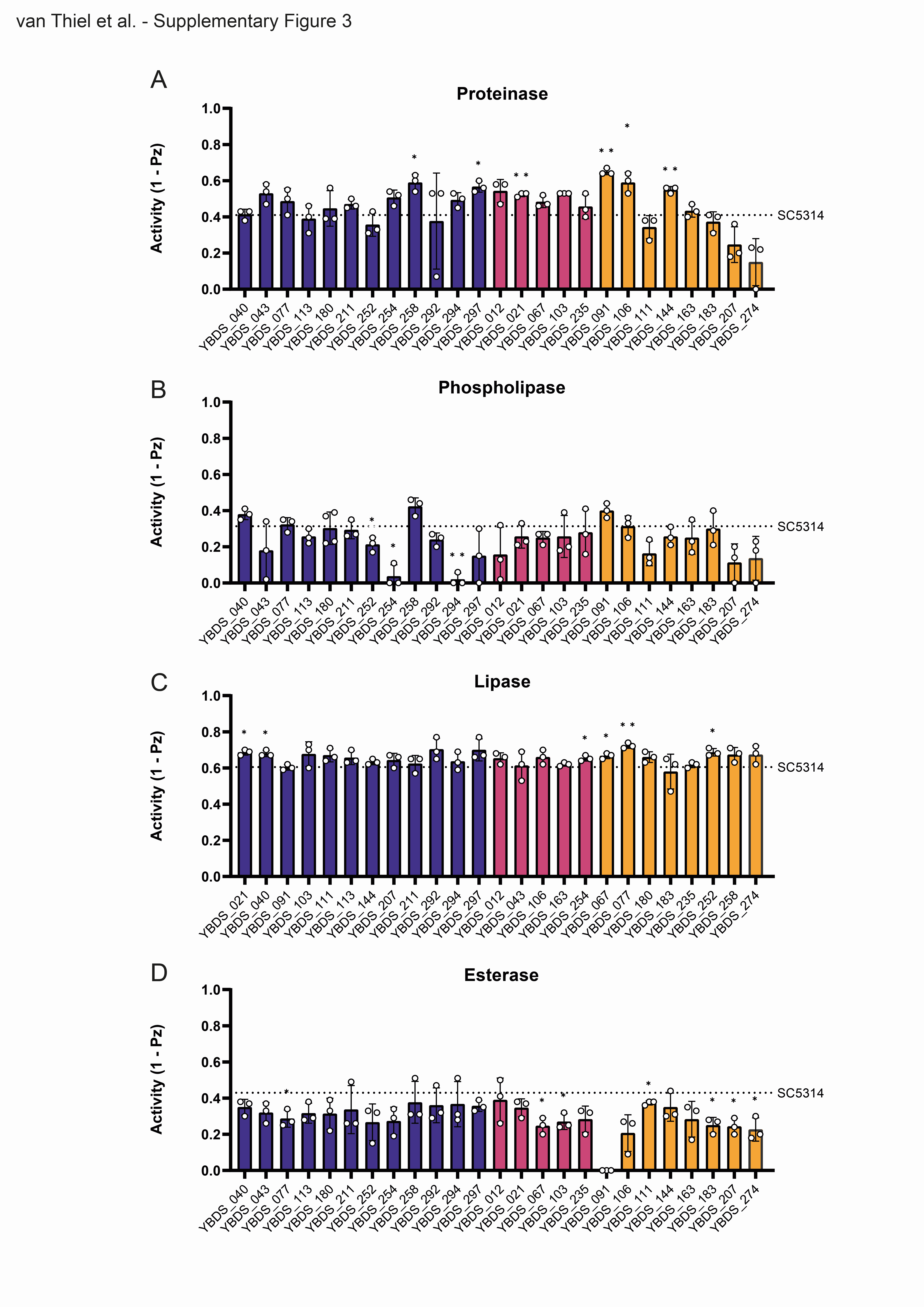

**Supplementary Figure 3 | Enzymatic activity of one feces-derived *Candida albicans* strain per individual.** A) Proteinase activity; B) Phospholipase activity; C) Lipase activity; D) Esterase activity. Dotted line represents average activity of reference strain SC5314. Statistical differences determined using one-sample t-test versus the relevant reference value. * p<0.05, * p<0.01.

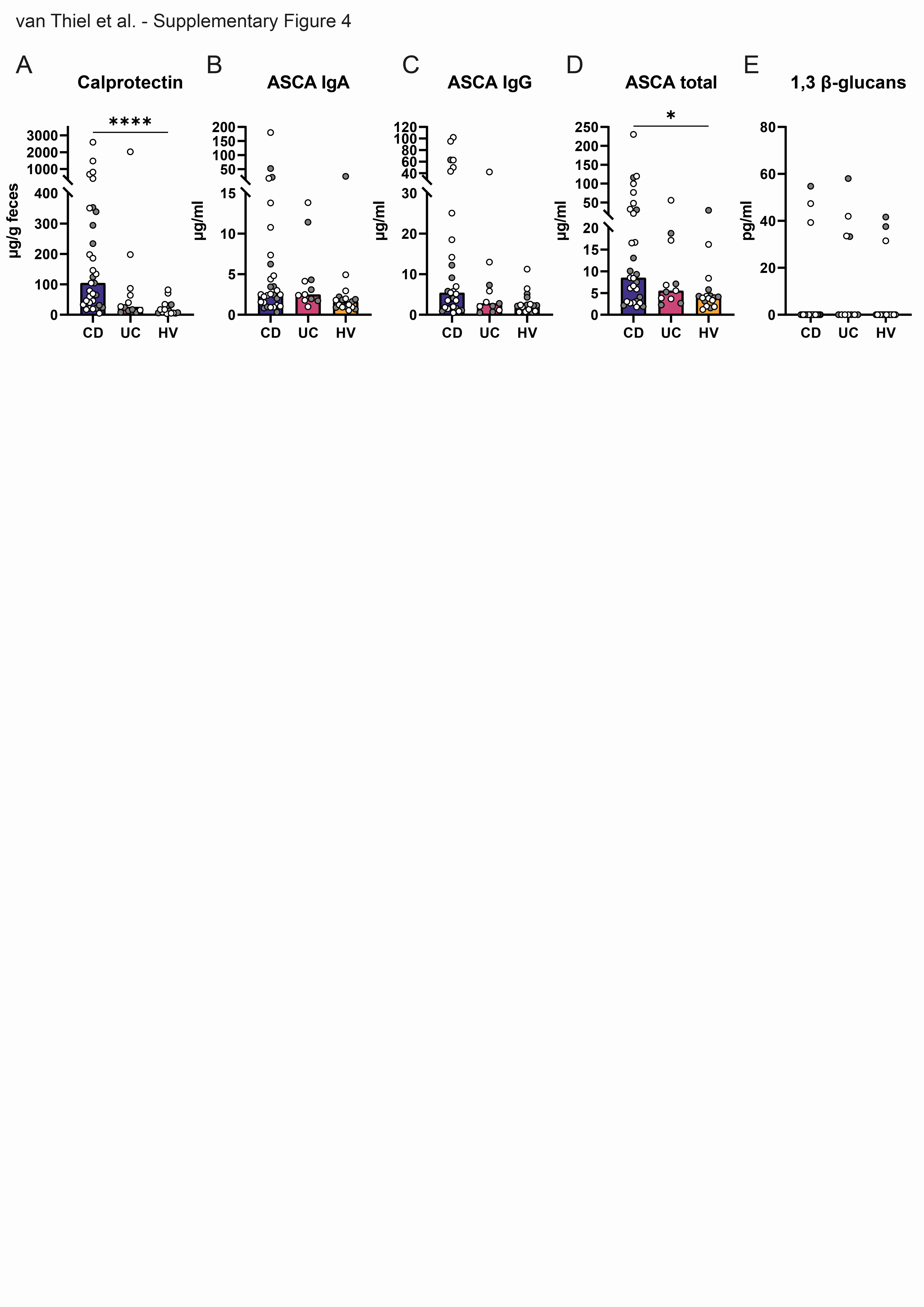

**Supplementary Figure 4 | Biomarker measurements in stool and serum of all patients.** A) Fecal calprotectin; B-D) *Anti-Saccharomyces cerevisiae* antibodies (ASCA) concentrations, divided in B) IgA, C) IgG, D) total concentrations; E) Serum 1,3-β-glucan levels. Gray datapoints refer to samples with corresponding positive *C. albicans* fecal cultures. Statistical differences determined using Kruskal-Wallis test. * p<0.05, **** p<0.001.
